# Characterization of Met25 as a Color Associated Genetic Marker in *Yarrowia lipolytica*

**DOI:** 10.1101/2020.03.30.017483

**Authors:** Harley Edwards, Zhiliang Yang, Peng Xu

**Affiliations:** Department of Chemical, Biochemical and Environmental Engineering, University of Maryland Baltimore County, Baltimore, MD 21250

**Keywords:** methionine biosynthesis, *Yarrowia lipolytica*, genetic marker, color associated screening, counter selection

## Abstract

*Yarrowia lipolytica* offers an ideal host for biosynthesis of high value natural products and oleochemicals through metabolic engineering despite being restricted to a limited number of selective markers, and counter-selection achieved primarily with *URA3*. In this work, we investigate *MET25*, a locus of sulfide housekeeping within the cell, to be exploited as a standard genetic marker. Divalent lead supplemented in media induces lead sulfide (PbS) aggregation in *MET25*-deficient cells such that deficient cells grow brown/black, and cells with functional copies of *MET25* grow white. Loss of *MET25* did not induce strict auxotrophic requirements for methionine in *Y*. *lipolytica*, indicating *MET25* deficiency could be rescued by alternative pathways. Plasmid and chromosomal-based complementation of *MET25* deficient cells on a double layer agar plate with nutrient gradients demonstrates delayed phenotype (white morphology) restoration, indicating post-transcriptional feedback regulation of methionine biosynthesis in this yeast. *MET25* deficient *Y. lipolytica* could be used as an efficient whole-cell lead sensor with detection limit as low as 10 ppm of lead in drinking water. We further tested whether *MET25* deficiency can be exploited to confer resistance to methyl-mercury through chemical neutralization and detoxification. Kinetic growth curves of wild type and *MET25*-deficient cells were obtained under varying concentrations of methylmercury and cellular toxicity to methyl mercury was calculated from the Hill equation. Our results indicate that methylmecury may not be used as the counter-selectable marker due to insignificant changes of growth fitness. This work demonstrates the utility of using *MET25* as a sensitive lead sensor and the challenges of using *MET25* as a counter-selectable genetic marker, as well as the complex regulation of methionine biosynthesis in *Y. lipolyitca*, which may shed lights for us to develop valuable biotechnological applications centering around the sulfur house-keeping metabolism of the nonconventional yeast.

## Introduction

With the limited number of auxotrophic markers in the oleaginous yeast *Yarrowia lipolytica*, establishing an additional counter selectable marker has potential to add significant value and versatility to the genetic toolbox with regards to engineering this host organism (Wong, Engel et al. 2017, Ma, Gu et al. 2020). *URA3* is the conventional selection marker used in a few yeast species including *Y. lipolytica* because it offers auxotrophic transformant screening as well as counterselection potential for marker removal, *via* 5’-FOA resistance (Fickers, Le Dall et al. 2003, Lv, Edwards et al. 2019). Great advancements in genome editing have been made in this host recently despite largely being restricted by this single genetic marker (Gao, Tong et al. 2016, Schwartz, Hussain et al. 2016, Gao, Tong et al. 2017, Schwartz, Shabbir-Hussain et al. 2017, Jang, Yu et al. 2018, Lv, Edwards et al. 2019). Counter selectivity is essential for performing iterative chromosomal gene integrations and ensuring that random integration of the selectable marker from the first round of genetic manipulations does not lead to false positives upon use of that marker in subsequent transformations (Lv, Edwards et al. 2019). The *LEU2* marker requires similar synthetically defined media, and an auxotrophic host strain (Le Dall, Nicaud et al. 1994, Larroude, Rossignol et al. 2018), but without any known methods offering counter selection. Hygromycin and the *hph* dominant marker give the advantages of antibiotic resistance selection (Holkenbrink, Dam et al. 2018, Wagner, Williams et al. 2018), largely decoupling cell death and fitness from essential nutrients, but includes disadvantages like dose-independent spontaneous resistance and still no counter selectivity (Otero and Gaillardin 1996). A mechanism in *S. cerevisiae* involving methionine provided counter selectivity, phenotypic indication and auxotrophy to methionine (Cost and Boeke 1996), and this had not yet been exploited in *Y. lipolytica.*

The *MET25* gene is found in *Y. lipolytica*, homologous to *MET15* found in *S. cerevisiae*, and various pathological species of *Candida*. *MET25* in *Y. lipolytica* encodes for O-acetyl homoserine sulfhydrylase (EC 2.5.1.49) with a length of 425 amino acids. This enzyme is found ubiquitously in various organisms (Yamagata 1976, Yamagata 1989), and is reported to be responsible for catalyzing numerous reactions involved in sulfur metabolism. Some of these proposed reactions are listed in Table 1, almost all of which are involved in metabolism of amino acids containing sulfur. This enzyme will catalyze an acetyl transfer reaction between methanethiol and O-acetyl-L-homoserine to produce methionine and acetate (reaction 1). Reactions 2 indicate the use of hydrogen sulfide (H_2_S) as a substrate in conjunction with other carbon backbone substrates. This indicates significant trans-activity with respect to various small carbon sources and the fixation/sequestration of hydrogen sulfide in yeast.

**Table 1.** Short list of proposed reactions catalyzed by O-Acetylhomoserine aminocarboxypropyltransferase. Reactions are found linked to EC 2.5.1.49 *via* KEGG database, and chemical reactions were drawn with ChemDoodle.

| Reactions | Proposed reactions catalyzed by O-Acetylhomoserine aminocarboxypropyltransferase (EC 2.5.1.49). |
| --- | --- |
| 1 | O-Acetyl-homoserine + Methanethiol → L-Methionine + Acetate |
| 2 | O-Acetyl-homoserine + Hydrogen sulfide → L-Homocysteine + Acetate |

The naming of the MET25 protein is somewhat confusing, as this locus has been studied by many other groups, in various hosts. *MET25*, *MET15*, and *MET17* are essentially synonymous in KEGG databases or NCBI catalogues. Since the discovery of *MET15* as a phenotypic locus, this same marker has been reported with methyl-mercury resistance (Singh and Sherman 1975, Ono, Ishii et al. 1991), for example, a color associated counter selectable marker was reported in *S. cerevisiae* (Cost and Boeke 1996), and this genetic tool has become a standard auxotrophic marker in *S. cerevisiae* strain BY4741 (Baker Brachmann, Davies et al. 1998, Sadowski, Su et al. 2007). Similarly, *MET15* has been utilized in *C. albicans* as a positive/negative color associated selection marker (Viaene, Tiels et al. 2000). *MET25* as a name seemingly came along when it was found that the promoter for O-acetyl homoserine sulfhydrylase in *S. cerevisiae* was highly tunable to methionine concentration *(Mumberg, Muller et al. 1994)*. The *MET25* promoter is used as a standard genetic part due to its tight tunability to methionine concentration*. S*pecifically, we characterized the function of the genetic sequence found by YALI0D25168g (of the GRYC database) in this work. There is no current report to establish the MET25 functionality in *Y. lipolytica.*

The recent development of genome-editing tools have made *Y. lipolytica* an ideal host for various applications, ranging from biofuel production (Xu, Qiao et al. 2016, Qiao, Wasylenko et al. 2017, Xu, Qiao et al. 2017), to natural product biosynthesis (Liu, Marsafari et al. 2019, Lv, Marsafari et al. 2019, Zhang, Zhang et al. 2019, Gu, Ma et al. 2020, Liu, Wang et al. 2020, Ma, Gu et al. 2020, Marsafari and Xu 2020) and commodity chemical manufacturing (Ledesma-Amaro, Dulermo et al. 2016, Cordova and Alper 2018, Gu, Ma et al. 2020). To further expand the genetic toolbox and understand the complex regulation of sulfur metabolism, we hypothesized that *MET25* could be utilized as a counter selectable color-associated genetic marker in *Y. lipolytica* and the complementation of MET25 will restore the cell phenotype. In this report, we used both homologous recombination and episomally expressed CRISPR-Cas12/cpf1 nuclease to disrupt *MET25*. We characterized the phenotype of *MET25*-deficient cells under the colorogenic media (soluble lead acetate) or counter-selectable media (methyl mercury). With plasmid-based or chromosome-based complementation of *MET25*, we also validated whether the phenotype could be restored. Based on a double-layer slanted agar with gradients of methionine, we inferred the post-transcriptional feedback regulatory mechanism underlying methionine biosynthesis. The development of *MET25* may further expand our ability to enable *Y. lipolytica* as an oleaginous yeast for various biosensing, bioremediation, bioproduction and biomedical applications.

## Results

### Chromosomal disruption of MET25 *via* homologous recombination

We first attempted to disrupt *MET25* via the conventional homologous recombination methods using a *URA3* disruption cassette. A black phenotype was observed on 37.5% (3/8) colonies picked from the CSM-Ura plate and spotted onto MLA (modified lead agar). (Supplementary Figure 1). This did not guarantee a true homogeneous population of *ΔMET25* mutants due to the possibility of mixed colonies from the CSM-Ura plate. This mixed population was confirmed visually by serial dilution and spreading on another MLA (modified lead agar) plate, where white and black colonies can be observed (Fig. 1A). This colony isolation was repeated (Supplementary Figure 2.A) until a true homogenous population of black colonies was observed, (Supplementary Figure 2.B)

**Fig. 1.**
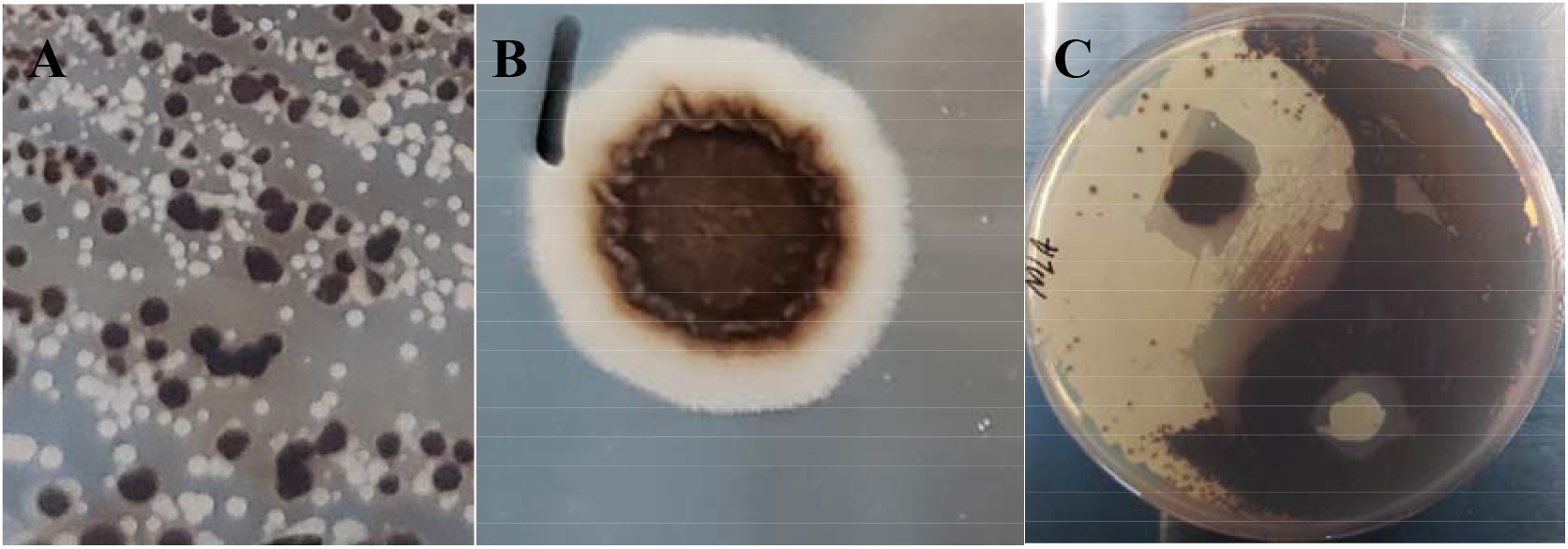
Visibly formed black colony of MET25 deficient cell. Phenotypic separation of white and black colonies after *MET25* deletion (A). Plasmid-complementation of *MET25* in the *MET25*-deficient cell leads to the formation of a radially distributed white ring (B). Yin-Yang art (C) of wild type (white colony) and *MET25* deficient cells (black colony) on MLA plate.

Results regarding methionine auxotrophic requirements vary from reports in other yeasts. Knocking out the *MET25* locus in *Y. lipolytica* did not produce a strict methionine auxotroph as *MET15* does in *S. cerevisiae* (Cost and Boeke 1996). Similarly, knockout of the appropriate homologue in *C. albicans* did not create an auxotroph (Viaene, Tiels et al. 2000). In order to test whether *MET25* is a strict methionine auxotrophic marker, colonies derived from Po1f, Po1f Δ*Met25*, and Po1fΔ*Met25* with pYLXP’-*Met25* were spotted on various media to confirm *MET25* deficiency and *MET25* restoration (Figure 2, and Supplementary Figure 3-6). The double dropout media, CSM-Leu-Ura plates, was used as control for the three genetic variants to ensure the correct genotype (supplementary figure 5). CSM-Met plates demonstrated to have little to no effective difference, indicating the existence of a methionine rescue pathway or contributions from media when spotting the plates. To rule out the pre-culture media effect, the cells were washed with PBS buffer and plated again onto CSM-Met (Supplementary Figure 3). We observed less growth in Δ*MET25* mutants, indicating clear cellular burden imposed on Δ*MET25* cells, although it did not induce a strict requirement for methionine. Cystathionine beta-lyase, *METC*, (YALI0D00605g) was hypothesized as the potential rescue pathway. The hypothesis was that by inhibiting the critical intermediate reaction between converting cystathionine to methionine, a methionine auxotrophic strain may be observed. The double knockout Po1fΔ*Met25*Δ*MetC*, also grew (data not shown) on methionine deficient media (CSM-Met), albeit with a diminished growth rate. We concluded that cystathionine beta-lyase does not appear to be a critical enzyme in the methionine rescue pathway responsible for conferring growth to our host without methionine.

**Fig. 2.**
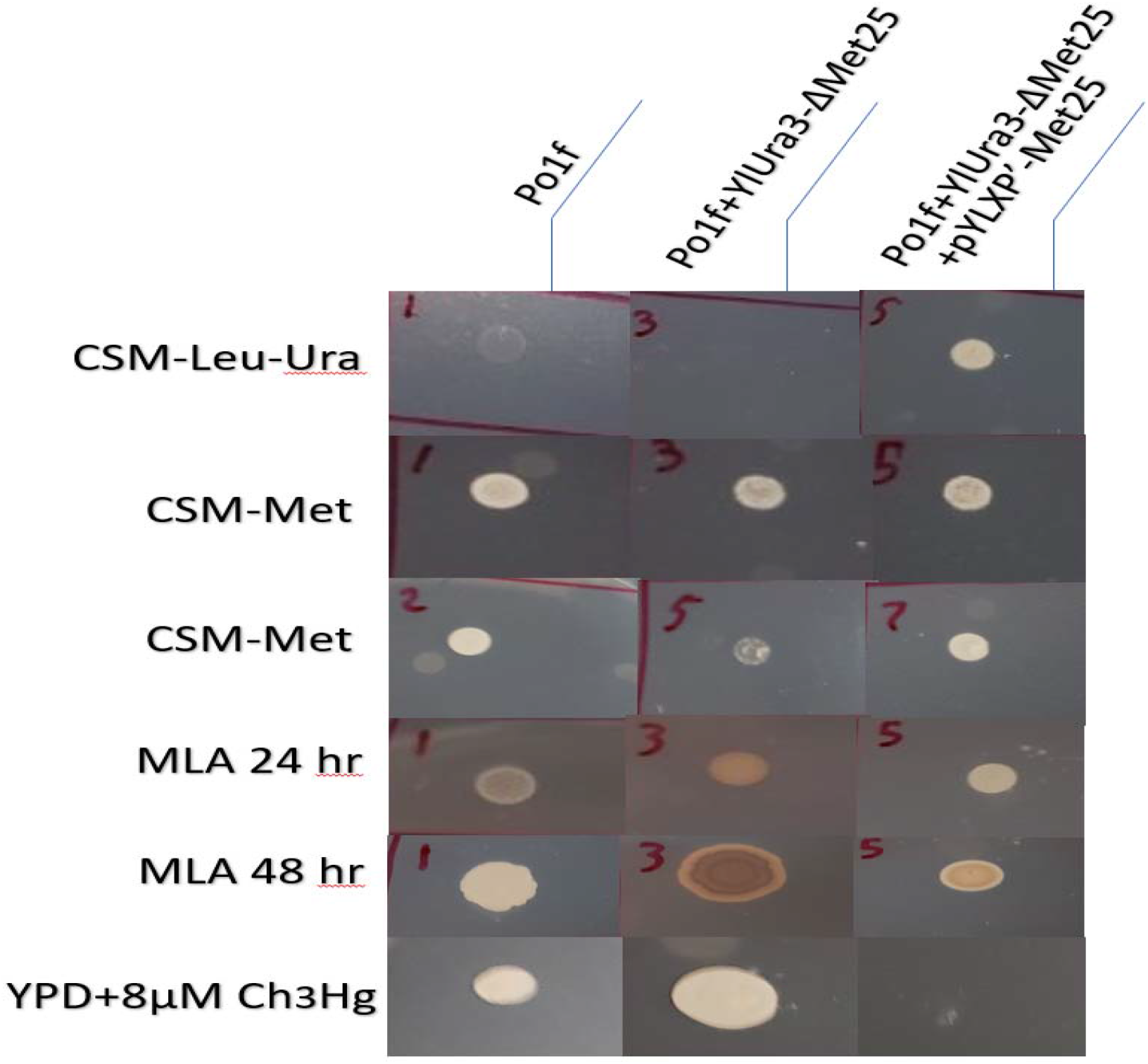
Collection of Selection Tests of MET25 deficient cell. This image contains the result various selective media assays. The columns left to right represent wildtype, mutant, and mutant containing restorative plasmid. The first row contains all strains growing on CSM-Leu-Ura. The second and third row contain all strains growing on CSM-Met, without and with cell washing, respectively. The fourth and fifth row demonstrates all strains growing on MLA, at 24 and 48 hours, respectively. The strains in the sixth row were grown on rich media containing 8mM methyl mercury. (The authors apologize for resolution lost in an attempt to demonstrate many pictures at once. Higher resolution images of the original plates are available in the supplementary information.)

### Genome-editing of *MET25 via* CRISPR-cas12/cpf1 nuclease

We also sought to disrupt *MET25 via* CRISPR-cas12/cpf1 in order to orthogonally demonstrate the same locus as a target for disruption with phenotypic indication. After transformation of Po1f with the dual expression plasmid pYLXP’-*AsCpf1*-*AsCrRNA*-*Met25*, colonies from CMS-Leu plates were then rescreened on MLA and grew for 72 hours for visible black sectoring to be observed in 1/16 samples. By the end of 7 days, 6 out of 16 sample had black sectors and after 14 days 11/16 had visible sectors (Figure 3). This indicates the genome-editing of *MET25* with CRISPR-Cpf1 depends on prolonged genome-targeting and cutting, as makes sense since the Cpf1 protein and associated RNA need time to be synthesized, folded, and located properly in vivo for their individual roles to be completed together. Phenotypically indicative sectors were colony isolated and *Met25* was sequencing verified to contain an indel knockout *(Yang, Edwards et al. 2020)*. The *MET25* locus provides an easy target for testing gene disruption efficiency with a phenotypic indication. *MET2*(YALI0E00836g), and *MET6(*YALI0E12683g) were all independently targeted with the same plasmid based Cpf1 gene disruption platform with success in achieving phenotypic selection (Yang, Edwards et al. 2020).

**Fig. 3.**
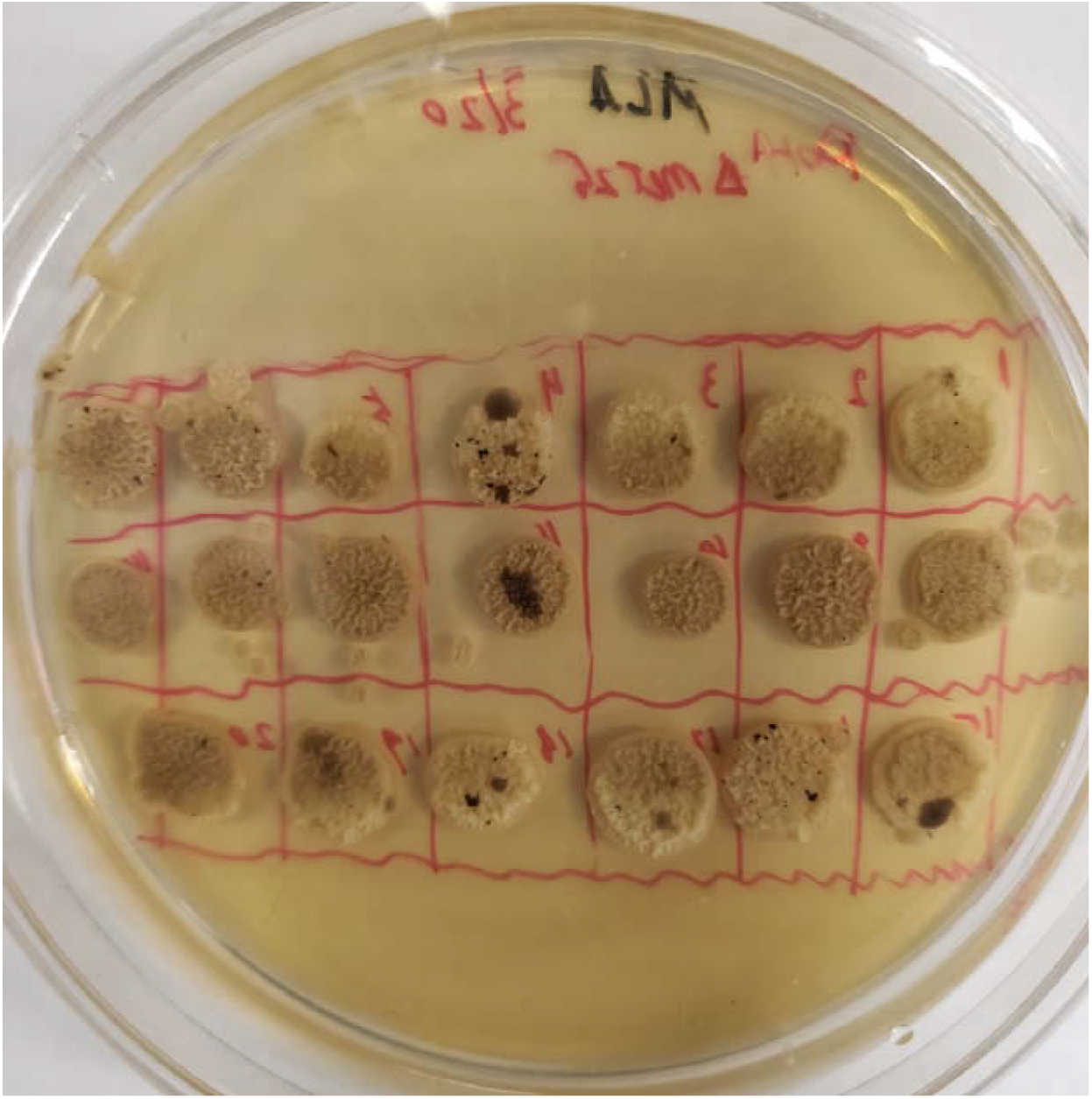
Visible Sectoring After Transient Disruption of MET25 via CRISPR-Cas12 and Targeted crRNA. Colonies from a successful transformation of Po1f with pYLXP’-AsCpf1-AsCrRNA-Met25 on CSM-Leu, subsequently spotted onto MLA. Performing colony isolation on any black sector results in successful indel knockout of MET25.

### Delayed phenotype restoration of *MET25* deficiency indicates orthogonal feedback regulation of methionine biosynthesis in *Y. lipolytica*

We next investigated whether we could restore the white colony phenotype by complementing *MET25* deficiency using either plasmid or chromosomal-based expression of *Met25*. Use of *MET25* alone in rich MLA media has proven difficult due to the lack of counter-selection pressure for negative transformants: almost all populations of transformants were black on the MLA plate, indicating either low transformation efficiency or negative transformants. To overcome this limitation, we next pursued a 2-step screening: we first screened the *LEU2* marker on CSM-leu plate to ensure expression of *MET25*, then the isolated colonies were replicated to MLA plate to validate phenotype restoration.

Episomal or chromosomal expression of *MET25* in *ΔMET25* cells proved difficult to restore the white colony phenotype, but the results of continued interest alluded to interesting regulation of methionine. Upon complementation with this *MET25* deficient host, the white phenotype was not observed initially (Fig. 1B). The cells grow in CSM-Leu, indicating the retention of the *LEU2* marker from the plasmid containing *MET25*. Re-streaking the colonies isolated from CSM-Leu also grew black on MLA, indicating no functional complementation of *MET25* from the plasmid or chromosomal-based *MET25* expression cassette, despite functional *LEU2* expression. After leaving the black transformant colonies in the incubator past 7 days, a white ring appeared radially surrounding a black core (Fig. 1B), as more generations grew. Colonies isolated from the white ring were re-streaked on MLA plates, demonstrating the similar patterns: a black core was surrounded by a white ring. MLA is rich media, and CSM-Leu/CSM-Met cannot stably contain the divalent lead without salting out so we cannot perform this phenotypic assay in minimal media. Since genetic knockout was validated by sequencing, we hypothesized that media contributions of methionine were causing the effect. Results from the first CSM-Met assays (Supplementary Figure 3), preliminarily indicated an extremely sensitive feedback mechanism to inhibit methionine synthesis by methionine despite *MET25* being expressed by an orthogonal promoter.

This hypothesis was further validated with the double-layer, slanted agar plate assay with a gradient concentration of methionine (Figure 4). The increasing size of the white ring was accompanied with a decreasing methionine concentration (Fig. 4), in both the plasmid and the chromosome-based complementation of the *MET25* gene. This assay confirmed that methionine was negatively autoregulating the functional expression of the *MET25* gene. For example, the critical outer diameter of the white ring was observed to be inversely proportional to the local methionine concentration (Fig. 4). In order to ensure it was methionine diffusion, a slanted agar plate was poured with MLA on bottom and CSM-Met on top and no coloration was observed (Fig. 4), indicating that Pb^2+^ diffusion was not causing the change of fitness of the cell.

**Fig. 4.**
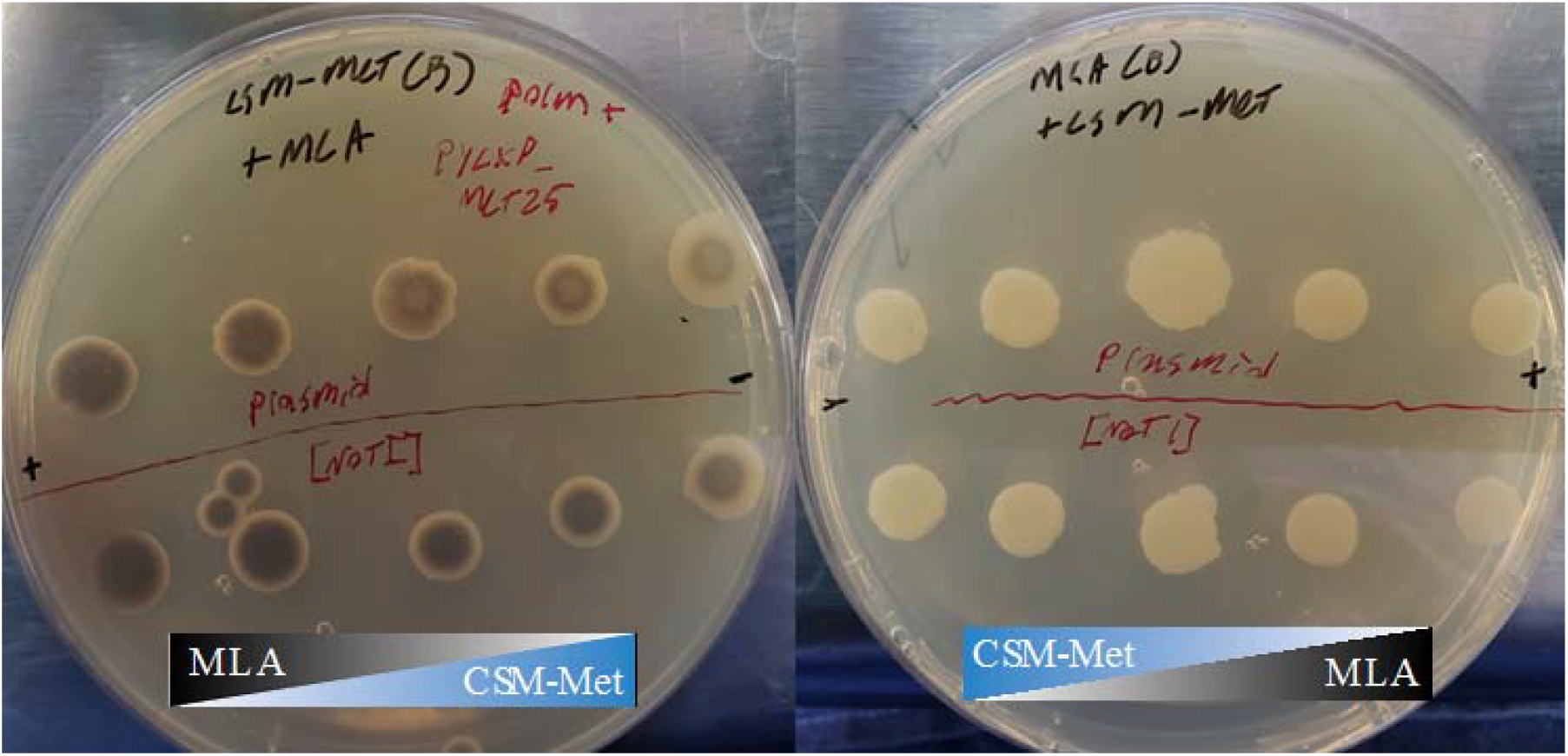
Dual-Media Slanted Agar Plates to detect MET25 phenotype restoration. On the left panel, CSM-Met on bottom, MLA on top, decreasing methionine availability from left to right. Phenotype restoration (white morphology) was observed to be increased with decreasing methionine availability, indicating the presence of methionine inhibits the expression of MET25 gene. On the right panel, MLA on bottom, CMS-Met on top, increasing methionine availability from left to right. All tested colonies remained white, indicating there is no lead diffusion from the bottom layer to the upper layer. Colonies with plasmid-based MET25 complementation are above the red equator line, colonies with chromosomally integrated MET25 (*via Not*1 linearization) are below the red equator line.

The white ring of the *MET25* complementation experiment suggests the methionine existing in the MLA media may produce feedback to inhibit the translation of *MET25* mRNA transcripts in *Y. lipolytica,* since it was orthogonally expressed via the *TEF2* promoter and *XPR2* terminator in our plasmid. The delayed phenotypic restoration in radially growing colonies indicated that there was sufficient amount of methionine inhibiting the expression of *MET25* in early growth stage, as a result, expression of *MET25* was not required, sulfur was not utilized, and the colonies remain black. At a later time when methionine was limited or depleted, expression of *MET25* was required, utilizing sulfur, and the new cells grew with their original white phenotype. It has been seen that there are multiple levels of regulation for *MET25* and methionine synthesizing pathways, including activating sequences of promoter regions, and post transcriptional interactions between methionine and the 5’ region of the mRNA transcript(Thomas, Cherest et al. 1989). In our case, the timing of the phenotypic shift is affected by methionine concentrations when expressed under an orthogonal promoter (*TEF2*) and terminator (*XPR2*), whether plasmid or chromosomally expressed. Our results allude to allosteric effects of methionine and Met25 the enzyme, co-repressor effects between methionine and an orthogonally expressed transcriptional factor, or post transcriptional activity between methionine and the *MET25* transcript.

### *MET25* deficient yeast as a whole cell senor to detect lead in potable water

Heavy metals in water have been linked with many diseases and have long been associated with neurodegenerative conditions in both kids and aged population. Since *MET25* deficiency leads to black pigmentation and form PbS precipitates, *MET25* deficient cells may be used as a whole-cell sensor to detect lead in potable waters. In order to gauge the applicability of this strain for microbially-based lead sensing purposes, the *ΔMET25* mutants were cultivated on MLA media containing 1000, 100, 10, and 0 mg/L (ppm), of soluble lead (II) nitrate, (Fig. 5). Yeast colonies and surrounding media were darkened in 100 ppm lead, by 24 hours, and significantly darker by 48 and 72 hours. Colonies gown on 10 ppm lead (II) MLA plates were visually darker after 72 hour’s cultivation (Fig. 5). These results indicate a high level of sensitivity could be achieved by a simple, microbial-based lead bioassay, reaching the detection limits of 10ppm (Fig. 5). Further genetic engineering may be needed to tune the sensitivity and dynamic range of this whole cell microbial probe even further. If the local sulfide concentration can be increased, or sulfate-related transporter genes could be manipulated, this organism could provide a scalable and effective, low cost, non-electric whole microbe sensor for heavy metals.

**Fig. 5.**
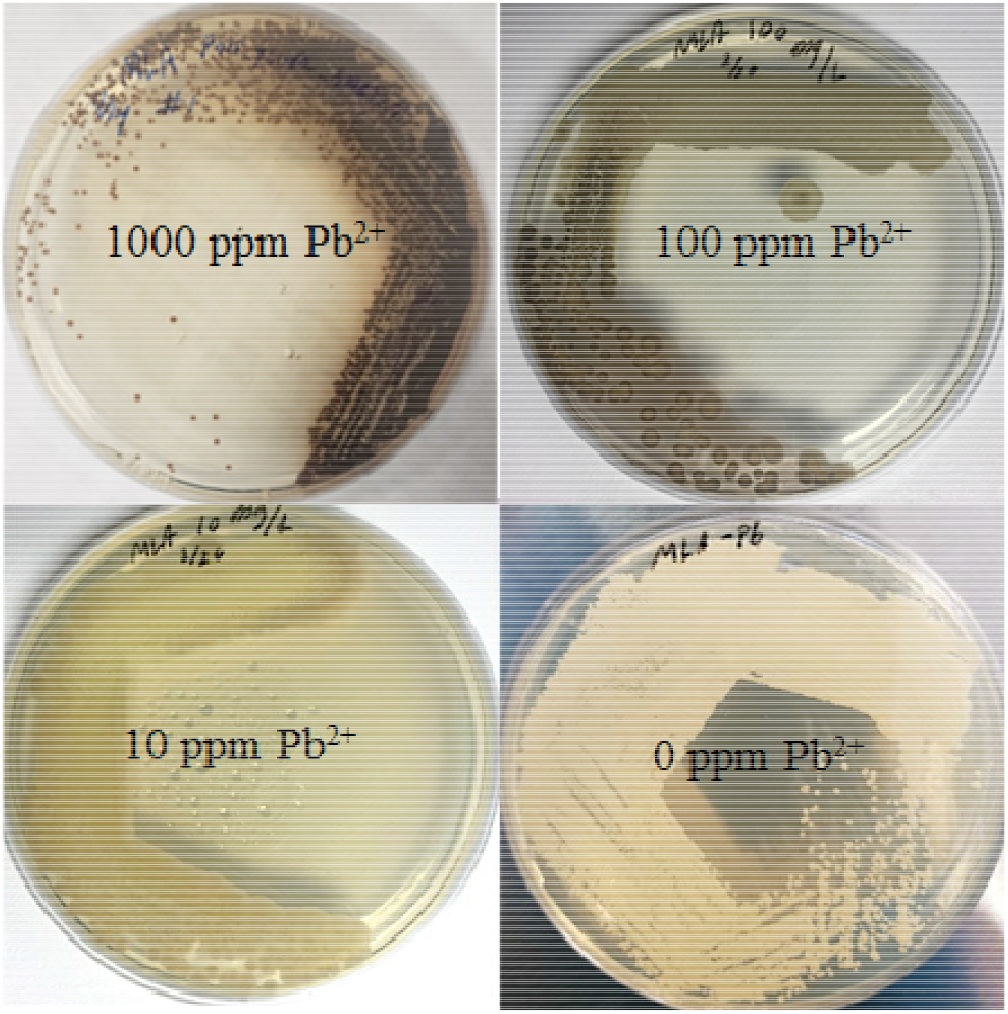
Phenotypic response of MET25 deficient strain to lead concentration ranging from 0 to 1000 ppm. MET25 deficient cells acting as biological probes for lead on MLA plates containing lead ranging from 0 ppm to 1000 ppm.

### Growth rates and counter-selection test toward methylmercury

We next attempted to determine the strength of the counter-selectivity toward methyl mercury in the *MET25* deficient strain. This specific growth rates for Po1f and Po1fΔ*Met25* were obtained by the slope of the linear regression of OD data versus time plotted on a logarithmic scale (equation No. 1). Then the specific growth rates were plotted against the level of methyl mercury to quantify the dose response relationship. Analyzing these specific growth rates in YPD at varying concentrations of methyl mercury in 96-well plates demonstrates a well fit with a Hill-type equation, governing the sigmoidal relationship between specific growth rate and methyl mercury concentration (Figure 6). The half inhibitory constants for Po1f and Po1f*ΔMet25* against methyl mercury were estimated at 0.746 μM and 1.383 μM respectively. As expected, with the deletion of the *MET25* gene, the mutant cells become more resistant to methyl mercury although by such a small margin, the practical applications (i.e. genetic selection with a <0.5 μM window of counter selectivity) are difficult to achieve.

**Fig. 6.**
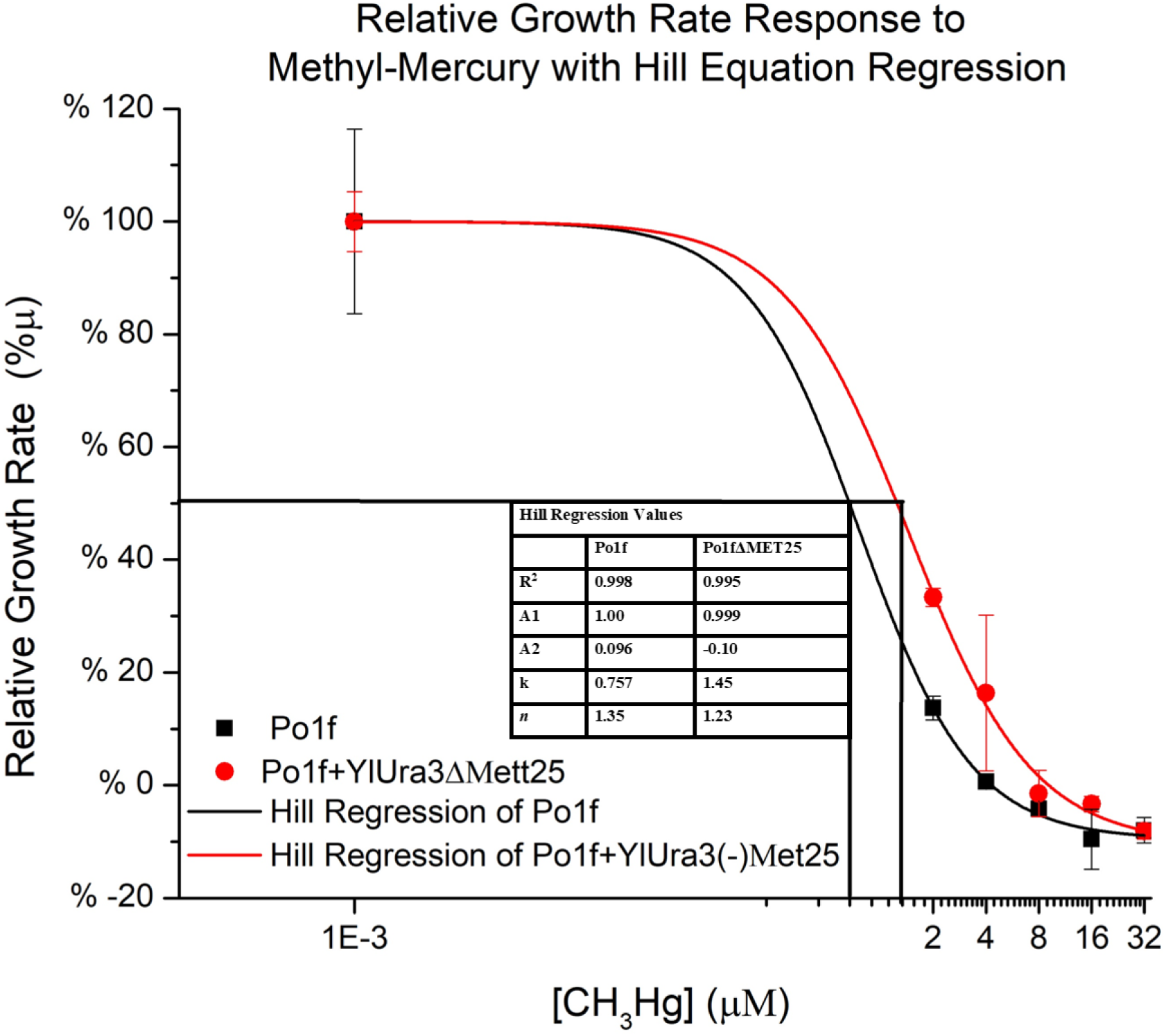
Relative Growth Rate Response to Methyl-Mercury. Relative specific growth rate in the presence of methyl mercury is observed to correlate strongly with a Hill type equation. Points indicate biological triplicates where the lines represent regression curves. Black graphics indicate wildtype and red indicate mutants lacking *MET25*.

## Discussion

Disruption of the *MET25* locus in *Y. lipolytica* via homologous recombination, or through targeted indel knockouts, certainly induces the formation of brown/black colonies in the presence of lead (II). The *ΔMET25* mutant generated visibly darker coloration to the naked eye on media containing as low as 10 ppm lead. These findings alone indicate a potential for biochemical engineers to develop a low cost, easy to use, microbial lead sensor for point of care applications in regions with polluted water. This was not the objective at the start of this experiment but the authors felt compelled to note this large dynamic range, and applicable sensitivity to a common heavy metal, after just one deletion. Further genetic engineering of microbial sulfur metabolism could increase lead detection sensitivity by increasing intracellular sulfide availability through limiting other metabolic steps that consume sulfide. The formation of lead (II) sulfide alludes to other possible sulfide and heavy metal chemistry, including the formation of color distinct cadmium sulfide or cadmium selenide.

CRISPR-Cpf1/Cas12-mediated genome editing targeting the *MET25, MET2*, or *MET6* loci was able to successfully induce black colony sectoring in Po1f, indicative of successful indel mutation or gene knockout (Yang, Edwards et al. 2020). These loci *MET25*, *MET2*, and *MET6* all provide similar behavior to target any of those genes for knockout, and a subsequent phenotypic screening. Knocking out any one of these *MET* genes individually does not induce an auxotrophic requirement for methionine, indicating that methionine can be biosynthesized via alternative route. Cystathionine β-lyase, EC.4.4.1.13, or *METC*, was knocked out, along with *MET25*, creating a double mutant which still was able to synthesize methionine to sustain growth. There is a methionine-adenine salvage cycle reported in plants and bacteria(Sauter, Moffatt et al. 2013), and we identified homologous proteins for the methionine salvage pathway in *Y. lipolytica*, pointing there for future genetic engineering targets to engineer a strict methionine auxotrophic strain. With the ubiquitous cellular requirement for methionine, considering it is also the start codon, it is likely advantageous for being redundant in pathways for housekeeping of enzymes responsible for methionine biosynthesis.

The chromosomal deletion of *MET25* induced resistance to the toxic chemical methyl-mercury and a Hill type relationship was demonstrated between the specific growth rate and the methyl-mercury concentration. The difference in half inhibitory constant (IC_50_) between the wildtype and mutant was about 0.6 μM, and this represents a challenge in exploiting this small window for counter selection. Since growth curves were in liquid media, those numbers were not helpful in determining a useful screening concentration of methyl-mercury on agar plates, and plate screening proved continuously difficult. Counter selection and phenotypic screening cannot be combined easily as the same intracellular sulfide affording a slight resistance to methyl mercury, is the same sulfide covalently reacting with lead to form a PbS black precipitate. If sulfide is utilized for color indication, it cannot also save the cells by neutralizing methyl mercury. Phenotypic screening and dropout media also could not be done together due to lead salts forming with nutrients in minimal media.

Most interestingly, even when *MET25* is expressed orthogonally via an independent promoter and terminator set (p*TEF2* and *XPR2*), there is still negative autoregulation of *MET25* activity due to local methionine availability. Further work should investigate differential gene expression under these conditions, and quest for the transcriptional factors (TFs) involved in this regulation, as well as perform calorimetric and energetic binding assay between the TFs, methionine, the *MET25* template DNA, and the *MET25* mRNA transcript. This may help us uncover a potentially novel feedback regulation mechanism in methionine biosynthesis apart from the commercialized *MET25* promoter. Considering the phylogenic age of the amino acid located at each and every start codon, there may be a robust and ubiquitous method of regulation like the tryptophan attenuation loop, or this could be a complex interaction network of transcription factors as is often not completely understood in eukaryotic signaling networks. Further investigation into the mechanism of feedback for methionine and *MET25, like quantitative RT-PCR to investigate the expression of critical genes in the pathway,* should be pursued for regulatory discovery. Further work should also standardize the phenotypic screening methods with the sulfur-housekeeping marker in *Y. lipolytica*, due to the very sensitive operating range of the methylmecury as a toxic selection agent.

## Conclusions

The *MET25* marker in *Y. lipolytica* can offer phenotypic screening of transformants quite easily on rich media supplemented with lead(II)acetate. Positive screening with *MET25* and solely lead has proven difficult as there is no negative selection pressure to limit growth of negative transformants and a methionine auxotrophic strain was not observed either. To overcome this limitation, counterselection is narrowly achieved via growth on rich media containing methyl-mercury. Methyl-mercury is toxic, and could be detoxified by sulfide buildup in the *ΔMET25* cells, due to the formation of mercuric sulfide and ethanol conferring toxic resistance to methyl mercury. The counter-selection window, at barely 0.5 micromolar in liquid media, makes this technique difficult to practically achieve. Both positive/negative screening, and counter selection can be done in rich media, although not in a single assay/plate due to synergistic effects of sulfide on both the mechanism of toxic resistance and the mechanism of phenotypic distinction.

*MET25* in our system was overexpressed via the *TEF2* promoter and *XPR2* terminator, but still, methionine-based autoregulation of this gene, observed in the slanted plate agar test, increased difficulty in consistent results with this strain. Counter selection was barely achieved and the narrow window of counter selectivity makes this technique difficult. Combined with the potential health hazards associated with mercury reagents and bioassays which use them, methyl-mercury selection is less than ideal. Further genetic engineering should work to build a methionine auxotroph, which would alleviate the necessity for methyl mercury counter selection, and should focus on increasing the rate and magnitude of this sulfide buildup to further leverage phenotypic and genetic selection traits. Utilizing these methods in rich media has potential to increase the rate at which transformants grow and are screened, which may significantly decrease the time spent on engineering this host. This selectable marker also facilitates better understanding of cellular regulation because of the visual indications of genetic events indicated by sectoring or delayed phenotype expression.

## Materials and methods

### Plasmids, strains and media

*Y.lipolytica* Po1f (ATCC MYA-2613, MATA ura3-302 leu2-270 xpr2-322 axp2-deltaNU49 XPR2:SUC2) was used as the host strain. *Escherichia coli* NEB5α was used for plasmid construction and proliferation. The YaliBrick plasmid pYLXP’ was used as the backbone to construct other plasmids (Wong, Engel et al. 2017). LB broth or agar plates containing ampicillin (100 mg/L) was routinely used for *E. coli* cultivation. YPD media consisting of 10 g/L yeast extract, 20 g/L peptone and 20 g/L glucose and complete synthetic media (CSM) omitting proper amino acids were used for yeast cultivation and transformation. MLA plates containing 3 g/L peptone, 5 g/L yeast extract, 0.2 g/L ammonium sulfate, 40 g/L glucose, 1 g/L lead nitrate and 20 g/L agar were used for visual selection of *met25* mutants. Lead nitrate was filter sterilized and added to autoclaved mixture of other components after cool down.

### Met25 Disruption via Homologous Recombination

All primers used in this work were listed in supplementary table 1. To construct a cassette for the deletion of *MET25*, primers met25upfw and met25uprv were used to PCR amplify an 800 bp fragment immediately upstream from the start codon of *MET25* using genomic Po1f DNA as template. This fragment was size verified via gel electrophoresis and purified using ZYMO Clean and Concentrator kits. Another 800 bp fragment immediately downstream from the stop codon was obtained using primers met25dwfw and met25dwrv. The gene *ylUra3* had previously been functionally cloned in our lab, into the plasmid pYLXP’, to create pYLXP’-*ylUra3*. That plasmid was linearized with *Sal*I and gel purified. The downstream 800 bp fragment was cloned into the *Sal*I digested vector backbone to yield pYLXP’-*ylUra3*-Met25DW, via Gibson assembly, transformed into *E. coli* and screened on LB agar plates.

Colonies were verified via colony PCR using xpr2_fw and met25dwrv. Positive colonies were inoculated into LB media containing ampicillin for overnight culture. Plasmid was purified using ZYMO Miniprep kits and sanger sequenced. The upstream 800 bp fragment was cloned into pYLXP’-*ylUra3*-Met25DW digested with *Cla*I to yield pYLXP’-*ylUra3*-Met25. The primers tef-rv and met25upfw were used for colony PCR to screen colonies for a 900 bp fragment. The sequencing-verified pYLXP’-*ylUra3*-*Met25* was used to PCR amplify a deletion cassette of *MET25* using primers met25cassfw and met25cassrv.

The *MET25* knockout cassette was transformed into wild type Po1f strain using hydroxyurea-based protocol to enhance homologous recombination(Tsakraklides, Brevnova et al. 2015),(Jang, Yu et al. 2018). Transformants were plated onto CSM-Ura plates. These preliminarily positive transformants were diluted into 10 μL sterile water and then used for selective media assays. These transformants were screened on MLA plates (Supplementary figure 1) and colonies which turned dark in color were picked to inoculate in CSM-Ura liquid media. That liquid culture was diluted and streaked on MLA to perform colony isolation. A single black colony was picked and inoculated into CSM-Ura liquid media, incubated 24 hrs, diluted, and re-plated to ensure that a true homogenous population of *ΔMET25* mutants was obtained. (Supplementary Figure 2)

### Plasmid and Chromosomal Complementation of Met25

*MET25* was PCR amplified out of genomic Po1f DNA using the primers met25fw and met25rv and cloned into vector pYLXP’ to yield pYLXP’-Met25. This plasmid was transformed into *ΔMET25* cells and plated onto CSM-Leu plates. Positive transformant were plated onto various selective medias (Supplementary Figures 3 to 6) for comparison of growth behavior and *MET25* expression in the wildtype Po1f, mutant Po1f+*ylUra3*-*Met25*, and mutant containing plasmid Po1f+*ylUra3*-*Met25* and pYLXP’-*Met25*. A graphic summarizing these plates can be observed in figure 2.

Growth assay in CSM-Met selective media were performed in order to test if the *MET25* knockout could confer auxotrophic requirements for methionine. Results were negative, as po1f-*ylUra3*-*ΔMET25* grew on CSM-Met plates when spotted from CSM-Ura liquid culture. The experiment was repeated to see if methionine in the liquid media was enabling growth. In this case cell cultures of each sample were grown for 2 days in appropriate selective media and centrifuged at 1800xg for 10 minutes to pellet cells. The supernatant was discarded, the pellet was resuspended in PBS and this entire wash was repeated to ensure no nutrient from the media is transferred with the cells that inoculate the selective media plates. These two plates are observed in Supplemental Figure 3.

To construct a strain with chromosomally integrated expression cassette of *MET25*, pYLXP’-*Met25* was linearized with NotI, a restriction site flanked by a chromosomal landing pad of complementary bases(Wong, Engel et al. 2017). The linearized plasmid was transformed into mutant cells and plated onto CSM-Leu plates. Colonies were inoculated into YPD liquid media for 72 hours and spread onto CSM-Leu plates. Single colonies from this plate were taken as chromosomal integrations of the plasmid.

### Met25 Disruption via CRISPR/Cpf1 Mediated Indel Mutation

In order to gauge the applicability of CRISPR/Cpf1 mediated, transient gene disruption of a phenotypic locus, and to orthogonally demonstrate the responsibility of the *MET25* gene in the lead(II) sulfide producing cells, a single plasmid was created containing a functional copy of *AsCpf1* and *AsCrRNA_Met25*. *AsCrRNA_Met25* is the gene enconding the guide RNA created analogous to the gRNA of the *CAN1* design(Wong, Engel et al. 2017). The construction of CRISPR-Cas12 plasmids used to knockout *MET25*, *MET2*, and *MET6* can be found in literature(Wong, Engel et al. 2017, Yang, Edwards et al. 2020). pYLXP’-*AsCpf1*-*AsCrRNA*-*Met25* was transformed into Po1f and plated onto CSM-Leu, incubated for 72 hours, and colonies were screened on MLA plates by picking single colonies into 10 μL sterile water, and spotting 3 μL on CSM-Leu, and 3 μL on MLA. These two plates were incubated for 48-72 hours (Supplementary Figure 4).

### MetC, and Met25 Double Knockout

A gene disruption cassette utilizing *ylUra3* marker was created and employed, using identical techniques as described in section 2.1, this time with 800 bp homologous arms designed to initiate homologous recombination in the *METC* (YALI0D00605g) locus. Colony PCR verification was performed using MetCupchk and tef-rv, as well as MetCdwnchk and xpr2-fw, for a 900 and 1000 bp fragment indicating successful integration, respectively. Positive colonies from colony PCR verification are now grown in CSM-Ura and designated as Po1f+*ylUra3*-*MetC*.

pYLXP’-*AsCpf1*-*AsCrRNA*-*Met25* was then transformed into Po1f+*ylUra3*-*MetC*. Transformants were grown on CSM-Leu for 72 hrs. Colonies were then picked, resuspended in 10 μL sterile water, and plated onto MLA plates. Once black sectoring was observed, this colony was chosen, diluted and spread onto MLA to isolate a single, black colony. This took three rounds of colony resuspension, dilution, plating, and isolation, before a homogeneous population was observed. This transformant was designated Po1f*ΔMet25*+*ylUra3*-*MetC*. This strain was assayed for an auxotrophic methionine requirement on CSM-Met plates.

### Selective Media and Differential Growth Test

All selective media agar plates use 2% agar. Dropout media CSM-Leu and CSM-Ura plates were used to selectively screen positive transformants. CSM-Met and CSM-Leu-Ura were employed to observe burden associated with different autotrophies, and to verify plasmid holding mutants respectively. Modified lead agar, MLA, plates were made with 1 g/L lead(II) nitrate(Van Leeuwen and Gottschling 2002), (Cost and Boeke 1996) YPD plates containing 4μM and 8μM methylmercury were used to investigate counter-selectivity. In order to establish a gradient of nutrients, agar plates were partially filled with CSM-Met, and the plates were tilted and allowed to cool such that the gel set diagonally. The plate was then poured the rest of the way with MLA media, ensuring to cover the minimal media entirely, and allowed to set again. This was repeated with MLA media on bottom and CSM-Met on top too. The results of the media assays with an established nutrient gradient are found in Figure 4. The authors apologize for resolution lost in attempt to demonstrate many pictures at once. Higher resolution images of the original plates are available in the supplementary information.

In order to more quantitatively observe response and gauge the utility of methylmercury as a counter-selection agent, specifically in Y. lipolytica, growth rates were measured in liquid media at varying concentrations of the methyl mercury. Po1f and Po1f+ylUra3-Met25 were inoculated in YPD and cultured for 24 hours. OD600 was normalized at 0.15 in the wells, and measurements were taken on a 96-well microwell-plate reader, during incubation at 30 degrees C with full shaking. OD was measured every 10 minutes for 8 hours. These readings were fit linearly on a logarithmic scale and the slope recorded as the specific growth rate. The specific growth rate was normalized with the maximum value to determine the relative growth rate labeled in percent of the maximum specific growth rate. These specific growth rates were plotted at various concentrations to visually establish a relationship for Po1f and Po1fΔMet25. Specific growth rates can be estimated by equation No. 1; and the inhibitory constant could be determined by the following Hill type equation (equation No. 2).

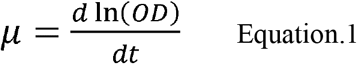

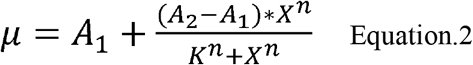

## Acknowledgements

This work was funded by the Bill & Melinda Gates Foundation under grant no. OPP1188443. as well as the National Science Foundation (Award Number 1805139). The authors also acknowledge the support from University of Maryland, Baltimore County, and the Department of Chemical, Biochemical, and Environmental Engineering.

## Author contributions

PX conceived and designed the topic. HE and ZY performed genetic engineering. HE wrote the manuscript. PX and ZY revised the manuscript.

