## Supplemental tables and figures for "Characterization of Met25 as a Color Associated Genetic Marker in *Yarrowia lipolytica*"

### Supplementary Table 1

**Primers used in this study**

| Primers | Sequence (5'-3') |
|---|---|
| MET25upfw | ggaattctcatgtttgacagcttatcCTAGCTTCTGAATCGCTACACAGTTC |
| MET25uprv | gtattccattatctacgaaaagcatcgatTGTGGATGTTAGTCTGAGGTGAGGTGG |
| MET25dwfw | ctagcgagacaataacggaggagtcgacTATCTTCATAACCACAACAGAGGACG |
| MET25dwrv | ctatgttacatccttttatcagacataGAGAGTGCTCACTGCGCTCTACAAG |
| MET25upchk | GTCACTATCCACGTGACCTAGCCACGC |
| MET25dwchk | GTTCCTAGACGCTACACGAAAGTAC |
| MET25cassfw | GCTTCTGAATCGCTACACAGTTCC |
| MET25cassrv | GAGTGCTCACTGCGCTCTACAAGG |
| MET25fw | cagcactttttgcagtactaaccgcagCCCTCGCACTTTGACACTCTGCAGCTC |
| MET25rv | ggggacaggccatggaactagtcggtaccCTAGTACACCGCCTTGAAAGCCTGC |
| ylMET25-crRR | GACGTTGATGCCGTAGGTCTTatctacaagagtagaaatta |
| Avr2_fw | gcatccctaaatttgatgaaagc |
| ylMET25-crRF | gtagatAAGACCTACGGCATCAACGTCggccggcatggtcccagcctc |
| Sal1_rv | GTTACATCCTTTTATCAGACATAGTC |
| Tef_fw | GGGTATAAAAGACCACCGTCCCC |
| Xpr2_rv | CCGTTGTAGGCAACAGCGTTGGG |
| MetC_Upfw | ggcgacgacggaattctcatgtttgacagcttatcACACCGAAATATCACTCTACAAG |

| | |
|---|---|
| MetC_UpRv | gtattccattatctacgaaaagcatcgatTTTTGTAGGGATCGCTCGTACTGTAG |
| MetC_DwFw | gctagcgagacaataacggaggagtcgacACCAATCCACAAAAATCCAATAAATT |
| MetC_DwRv | cttgcctatgttacatccttttatcagacataGTATATATGGGACCTCTCCCCCG |
| MetC_CassFw | CACCGAAATATCACTCTACAAGTATAG |
| MetC_CassRv | ATATATGGGACCTCTCCCCCGTTC |
| MetC_upChk | CCATTCACATTGAACAGTGTACAG |
| MetC_DwChk | CAATCCTCAATCCTAAACAAACC |
| ylMET2(A)-crRF | gtagatCTGGTCCGAGACCAGTCCCGAggccggcatggtcccagcctc |
| ylMET2(A)-crRR | GACGTTGATGCCGTAGGTCTTatctacaagagtagaaatta |
| ylMET2(B)-crRF | gtagatCCACGCTCTTACCGGTTCCGCggccggcatggtcccagcctc |
| ylMET2(B)-crRR | TCGGGACTGGTCTCGGACCAGatctacaagagtagaaatta |
| ylMET2_checkFW | GCAAGGAGAAGCCGAACCGTAGGG |
| ylMET2_checkRV | CACATCTTTTCATCTTCTTCTACAACCA |

**Supplementary Table S2.** Plasmids and Strain genotype used in this study.

| Strains or plasmids | Description | Reference |
|---|---|---|
| **Strains** | | |
| *E. coli* NEB 5α | *fhuA2* Δ(*argF-lacZ*)*U169 phoA glnV44 Φ80 Δ(lacZ)M15 gyrA96 recA1 relA1 endA1 thi-1 hsdR17* | New England Biolabs |
| *Y. lipolytica* po1g | MATa, *leu2*-270, *ura3*-302::URA3, *xpr2*-3 | Lab stock |
| *Y. lipolytica* po1f | MATa *ura3*-302 *leu2*-270 *xpr2*-322 *axp2*-deltaNU49 *XPR2*::*SUC2* | Lab stock |
| *Y. lipolytica* | po1f+*ylUra3-Met25* | This work |
| *Y. lipolytica* | po1f+*ylUra3-MetC* | This work |
| *Y. lipolytica* | po1fΔ*Met25+ylUra3-MetC* | This work |
| *Y. lipolytica* | po1fΔ*Met25* | This work |
| *Y. lipolytica* | po1fΔ*Met6* | This work |
| *Y. lipolytica* | po1fΔ*Met2* | This work |
| **Plasmids** | | |
| pYLXP' | Cloning vector | Lab stock |
| pYLXP'+ *-Met25* | MET25 restoration plasmid | This work |
| pYLXP'+*ylUra3-Met25* | Homologous recombination MET25 knockout cassette construction plasmid | This work |
| pYLXP'-*ylUra3-MetC* | Homologous recombination METC knockout cassette construction plasmid | This work |
| pYLXP'-*AsCpf1-AsCrRNA-Met25-* | CRISPR/Cas12 nuclease expression plasmid with gRNA targeting MET25 for indel knockout | This work |
| pYLXP'-*AsCpf1-AsCrRNA-Met2-* | CRISPR/Cas12 nuclease expression plasmid with gRNA targeting MET2 for indel knockout | This work |
| pYLXP'-*AsCpf1-AsCrRNA-Met6-* | CRISPR/Cas12 nuclease expression plasmid with gRNA targeting MET6 for indel knockout | This work |

**Supplementary Figure 1**

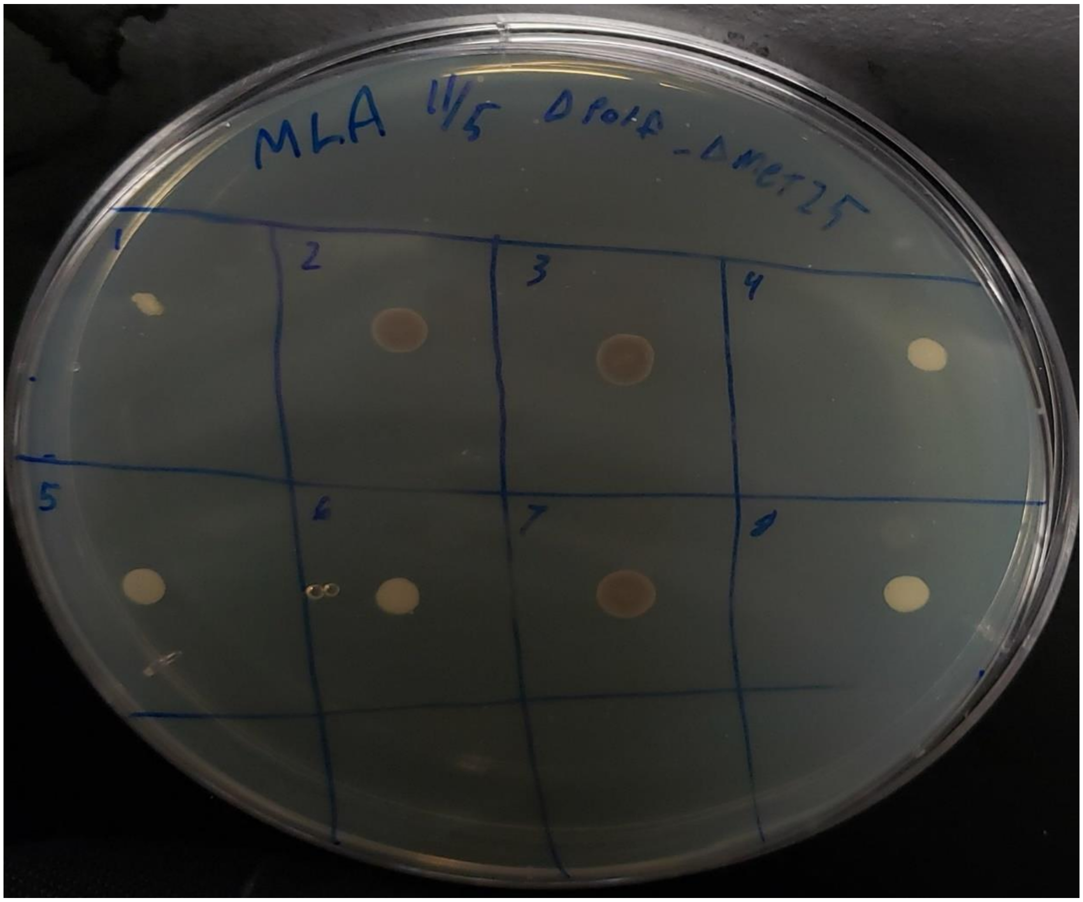


**Supplementary Figure 1.** Preliminarily positive transformants on MLA media, selected immediately after cassette transformation and growth on CSM-Ura plates.

**Supplementary Figure 2**

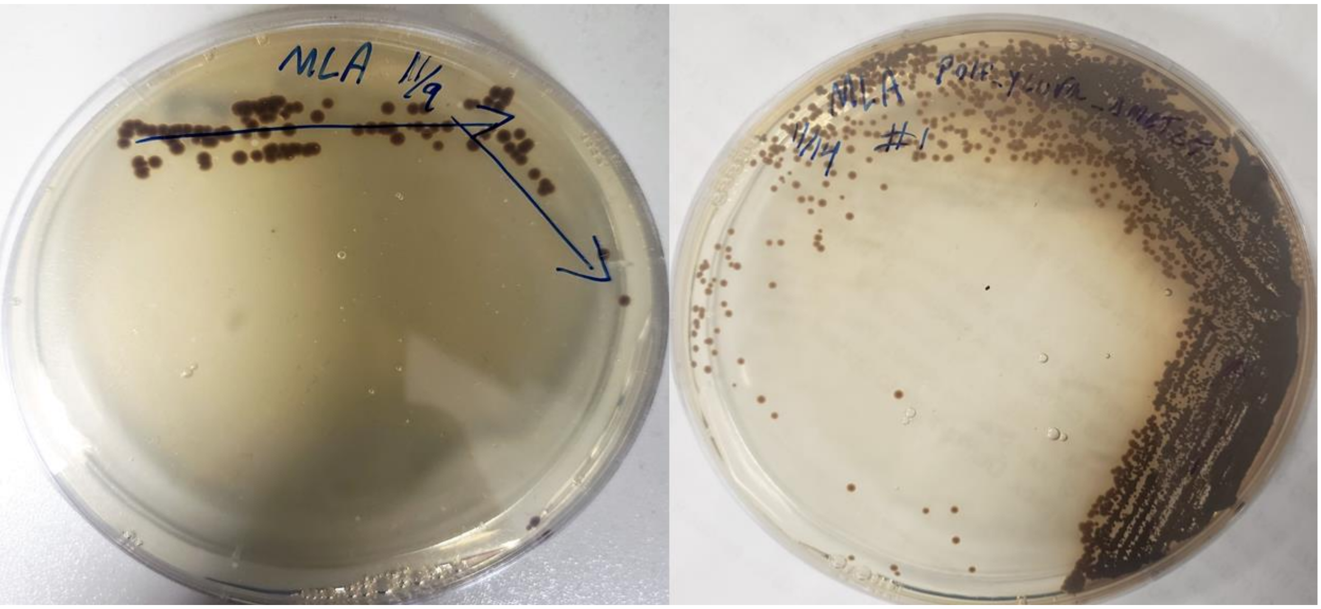


**Supplementary Figure 2.** 2.A On the left is the result of growth from a culture of positive ΔMET25 mutants from Figure 1, after MLA and subsequent 24 hour incubation in CSM-Ura liquid media. Two white colonies can be observed indicating cells with functional copies of MET25. 2.B was the result of a single colony from the plate in 2.A, inoculated into CSM-Ura, cultured for 24 hours, diluted, and re-plated on MLA.

**Supplementary Figure 3**

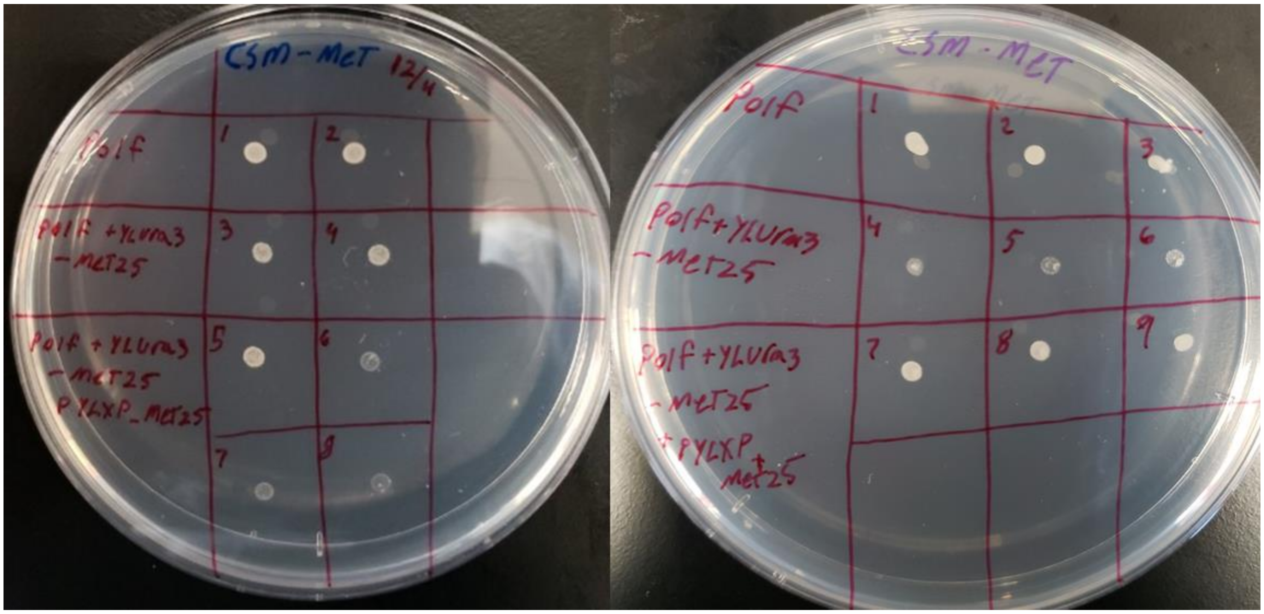


**Supplementary Figure 3.** Top to bottom there is the wildtype, ΔMET25 mutant, and the mutant with the restorative plasmid. On the left are cells spotted directly from culture solution. On the right, the cells were first pelleted, and resuspended in PBS, twice.

**Supplementary Figure 4**

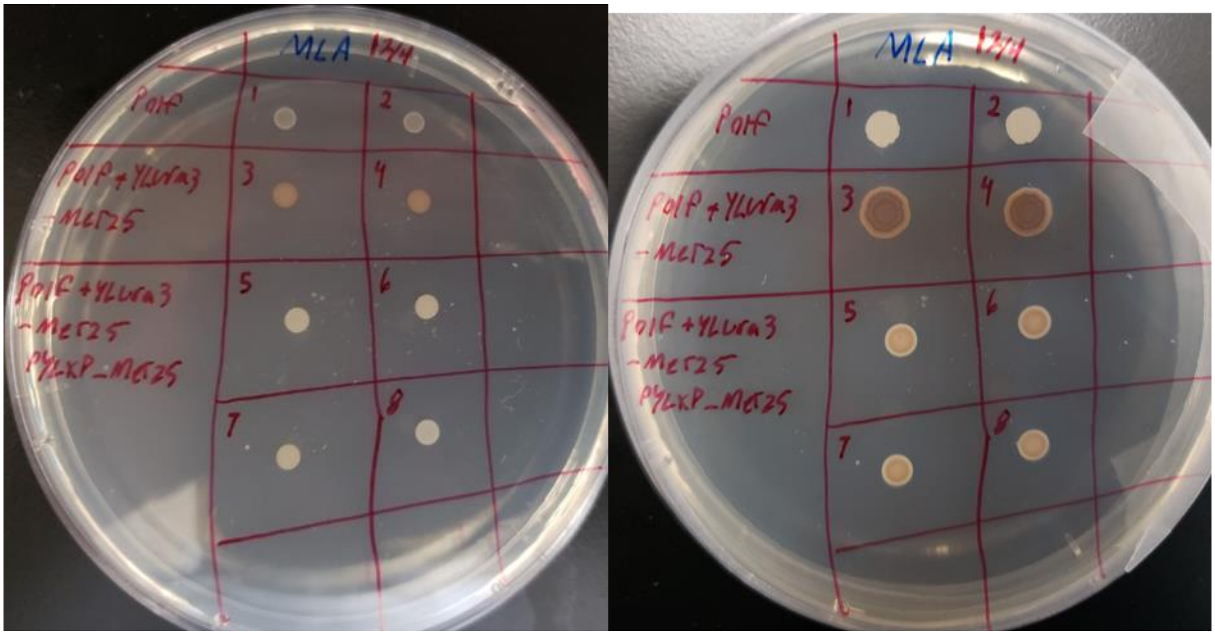


**Supplementary Figure 4.** Here are two images of growth on rich media containing divalent lead. In the top section the wildtype is white. In the second row, the MET25 mutant, and on the third and fourth row, the plasmid restoration of MET25 activity. On the left is 24 hours after spotting on the plate, on the right 48 hours.

**Supplementary Figure 5**

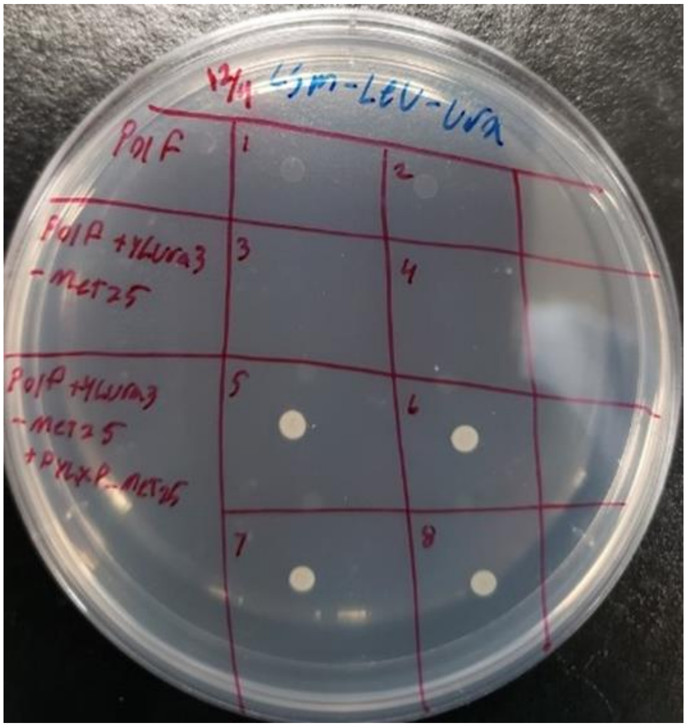


**Supplementary Figure 5.** This is the control plate to ensure both markers were functionally expressed in our testing. The first row is the wildtype, second is the MET25 mutant, third is the mutant with the restorative plasmid.

**Supplementary Figure 6**

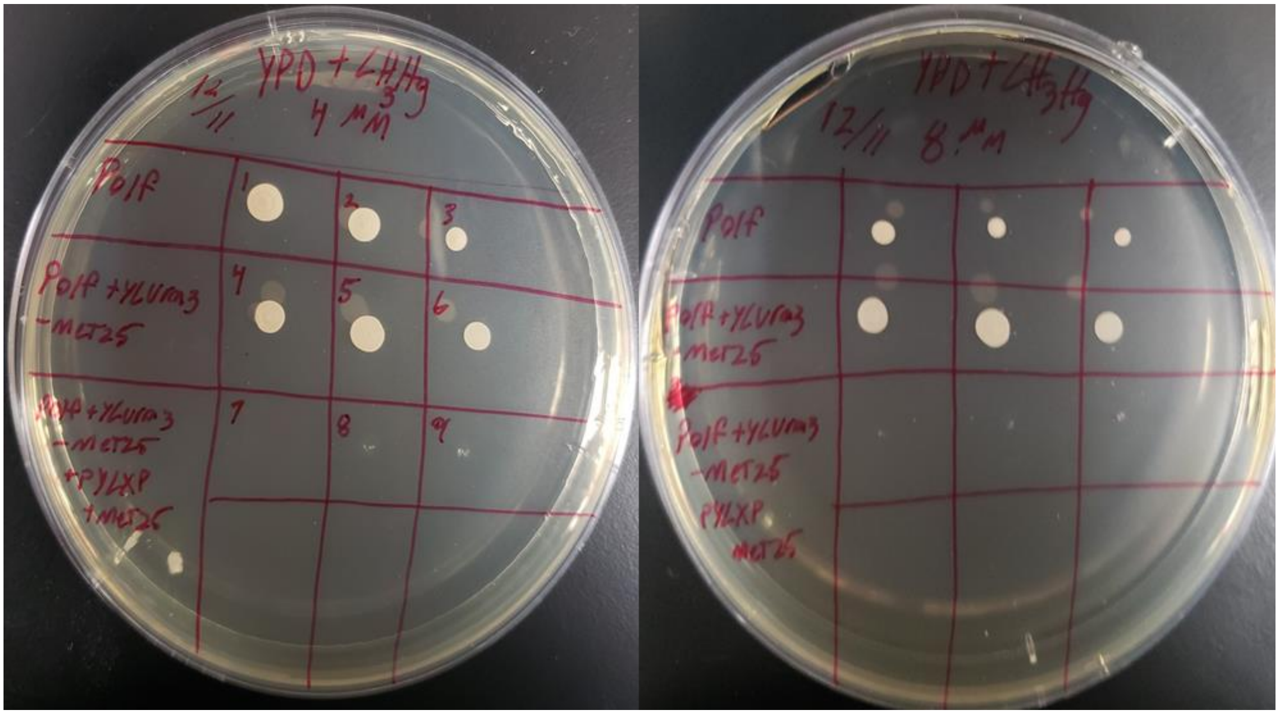


**Supplementary Figure 6.** On the left, the concentration is 4μM methyl mercury in YPD, on the right, 8μM. Under 8μM, we can see a slight advantage afforded to the MET25 mutant cells.
